# CellWalker integrates single-cell and bulk data to resolve regulatory elements across cell types in complex tissues

**DOI:** 10.1101/847657

**Authors:** Pawel F. Przytycki, Katherine S. Pollard

## Abstract

Single-cell and bulk genomics assays have complementary strengths and weaknesses, and alone neither strategy can fully capture regulatory elements across the diversity of cells in complex tissues. We present CellWalker, a method that integrates single-cell open chromatin (scATAC-seq) data with gene expression (RNA-seq) and other data types using a network model that simultaneously improves cell labeling in noisy scATAC-seq and annotates cell-type specific regulatory elements in bulk data. We demonstrate CellWalker’s robustness to sparse annotations and noise using simulations and combined RNA-seq and ATAC-seq in individual cells. We then apply CellWalker to the developing brain. We identify cells transitioning between transcriptional states, resolve enhancers to specific cell types, and observe that autism and other neurological traits can be mapped to specific cell types through their enhancers.

---

Gene regulatory elements are critical determinants of tissue and cell-type specific gene expression^1,2^. Annotation of putative enhancers, promoters, and insulators has rapidly improved through large-scale projects such as ENCODE^3^, PsychENCODE^4^, B2B^5^, and Roadmap Epigenomics^6^. However, both predictions and validations of regulatory elements have been made largely in cell lines or bulk tissues lacking anatomical and cellular specificity^7^. Bulk measurements miss regulatory elements specific to one cell type, especially minority ones^8^. This lack of specificity limits our ability to determine how genes are differentially regulated across cells and to discover the molecular and cellular mechanisms through which regulatory variants affect phenotypes.

Single-cell genomics is an exciting avenue to overcoming limitations of bulk tissue studies^8,9^. However, these technologies struggle with low-resolution measurements featuring high rates of dropout and few reads per cell^8,9^. Many methods have been developed to address these problems in single-cell expression data (scRNA-seq)^8,9^. However, these strategies generally fail on scATAC-seq data because there are fewer reads per cell, and the portion of the genome being sequenced is typically much larger than the transcriptome^10^. Consequently, scATAC-seq has much lower coverage and worse signal-to-noise than scRNA-seq.

Several scATAC-seq analysis methods have been developed to increase the number of informative reads used per cell. These include CICERO^11^, which aggregates reads from peaks that are co-accessible with gene promoters to emulate gene-focused scRNA-seq data, and SnapATAC^12^, which computes cell similarity based on genome-wide binning of reads. Other methods search for informative reads based on known or predicted regulatory regions^13,14^. However, these approaches often miss rare but known cell types^10^. Other methods attempt to detect cell types in scATAC-seq data by either mapping the data into the same low-dimensional space as scRNA-seq data or by labeling cells in scATAC-seq to known cell-type expression profiles^15,16^. While these provide a promising avenue towards adding labels to clusters of cells observed in scATAC-seq data, they do not help to increase the resolution of cell type detection.

We present CellWalker, a generalizable network model that improves the resolution of cell populations in scATAC-seq data, determines cell label similarity, and generates cell-type specific labels for bulk data by integrating information from scRNA-seq and a variety of bulk data. These labels can be generated concurrently from the same tissue, but could also be from cell lines, sorted cells, or related tissues. Our method goes beyond co-embedding or directly labeling cells with this prior knowledge about cell types, instead propagating cell-type signatures over a network of cells and cell types so that they are weighted with evidence of cell types in scATAC-seq. Diffusion through this network allows labeling information to indirectly influence cells with similar genome-wide open chromatin profiles even if they could not be initially labeled. A major benefit of our model is that it allows us to compute the level of influence of each label and cell on every other label and cell, thus providing an avenue for additional inferences. These include deconvoluting bulk measurements and assessing their relevance to specific cell populations, as well as quantifying similarity between known cell types in the tissue where scATAC-seq was performed.

The developing human brain presents a complex landscape of cell types each with unique regulatory programs^17–19^. Using CellWalker we mapped cell types derived from scRNA-seq data to a large set of scATAC-seq data. The derived influence matrix made it possible to examine changes in regulation across neuronal development and map enhancers to specific cell types. Using this cell-type specific atlas of enhancers we found that autism spectrum disorder (ASD) genes are enriched for enhancers specifically active in inhibitory interneurons, while developmental delay genes are enriched for enhancers specifically active in radial glia. The ability to map psychiatric traits to cell types is a crucial step towards understanding the mechanisms through which disease develops and responds to treatment. As more large-scale single cell studies are released, generalizable methods such as CellWalker will be fundamental toward integrating them with existing bulk data to increase our understanding of cell-type specific regulatory programs.

## Results

### Overview of method

CellWalker resolves cell types and differentially accessible regions in scATAC-seq data by integrating information from scRNA-seq and bulk data. This integration relies on building a combined network featuring nodes representing cells in scATAC and nodes for external labeling data, e.g. cell types derived from scRNA-seq data (Figure 1a). Briefly, cells from scATAC-seq are nodes in the network, and edges between them encode information about cell similarity. A second set of nodes represents labeling datasets connected to cell nodes by edges that encode the similarity between a label and a cell. Using a graph diffusion implemented via a random walk with restarts, CellWalker computes a global influence matrix that defines the influence of every cell and label on every other cell and label (Figure 1b). In this matrix, each column represents where walks starting from that node end. Different portions of this matrix can be used to map information between and within domains: cell-to-cell for clustering cells, label-to-label for exploring label similarity, label-to-cell for cell type labeling, and cell-to-label for distributing bulk signatures to labels.

**Figure 1.**
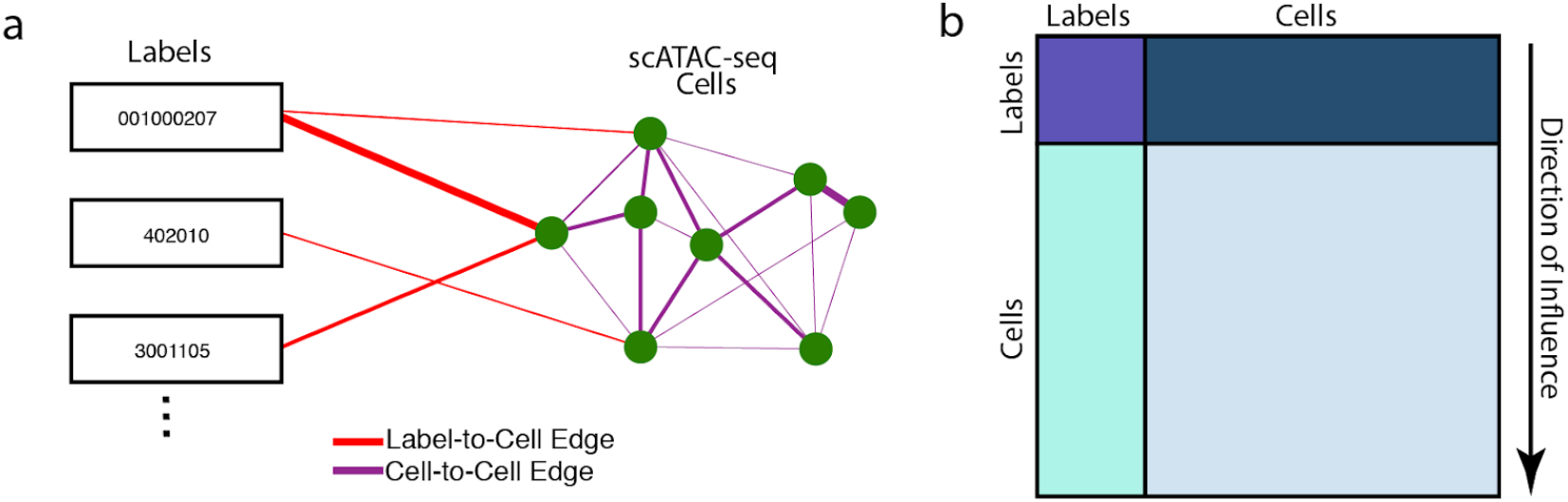
Overview of CellWalker. **a.** Cells (circles) are connected based on similarity of their scATAC-seq profiles (purple edges). The network is extended using external data labels (e.g. expression levels of marker genes for a cell type, rectangles). These labels are connected to cells based on how well they correspond to each cell (e.g. fraction of cell’s promoters accessible in marker genes, red edges). Information is propagated across the combined network using global diffusion. **b.** Diffusion results in an influence matrix that describes the information passed from each label and cell to every other label and cell. Label-to-label influence (purple) encodes label similarity, label-to-cell influence (teal) encodes which labels best describe each cell, cell-label influence (blue) can be used to map information encoded in accessibility back to labels, and cell-to-cell influence represents a label-influenced clustering of cells (light blue).

### Method validation and evaluation

To assess the ability of CellWalker to distribute labeling information across cells we first tested it on simulated data. For each simulated scenario we tuned a single label edge weight parameter defining the ratio of label-to-cell edges versus cell-to-cell edges (Figure 2a). This parameter represents a trade-off between information about cell labels versus scATAC-seq-based cell similarities. When edge weight is low the output is similar to de-novo cell clustering using only scATAC-seq, and when it is high the output converges towards directly assigning labels to each cell. To quantify performance, we developed a measure called cell homogeneity which is computed directly from the influence matrix as the median ratio of information between cells within the same cell type to information between cells of different cell types. A higher cell homogeneity indicates a greater ability to differentiate between different cell types. We found that as few as 10% of cells being labeled is sufficient for CellWalker to improve cell labeling as measured by cell homogeneity and that there is no further improvement after ~30% of cells are labeled (Figure 2b). As expected, performance degrades as more cells are initially mislabeled and improves when cells of different types are more distinct from each other in the scATAC-seq data (Figure 2c and Supplemental Figure 1a). Furthermore, CellWalker performs well with noisy data, even when up to 50% of reads are dropped or random reads are added (Supplemental Figure 1b). Finally, we observe that CellWalker is able to distribute labels to novel cell populations (Supplemental Figure 1c). Importantly, CellWalker achieves this efficiently in terms of both time and computer memory usage (Supplemental Figure 2). These results establish the network diffusion strategy implemented in CellWalker as a robust approach to integrate scATAC-seq with scRNA-seq or other labeling data.

**Figure 2.**
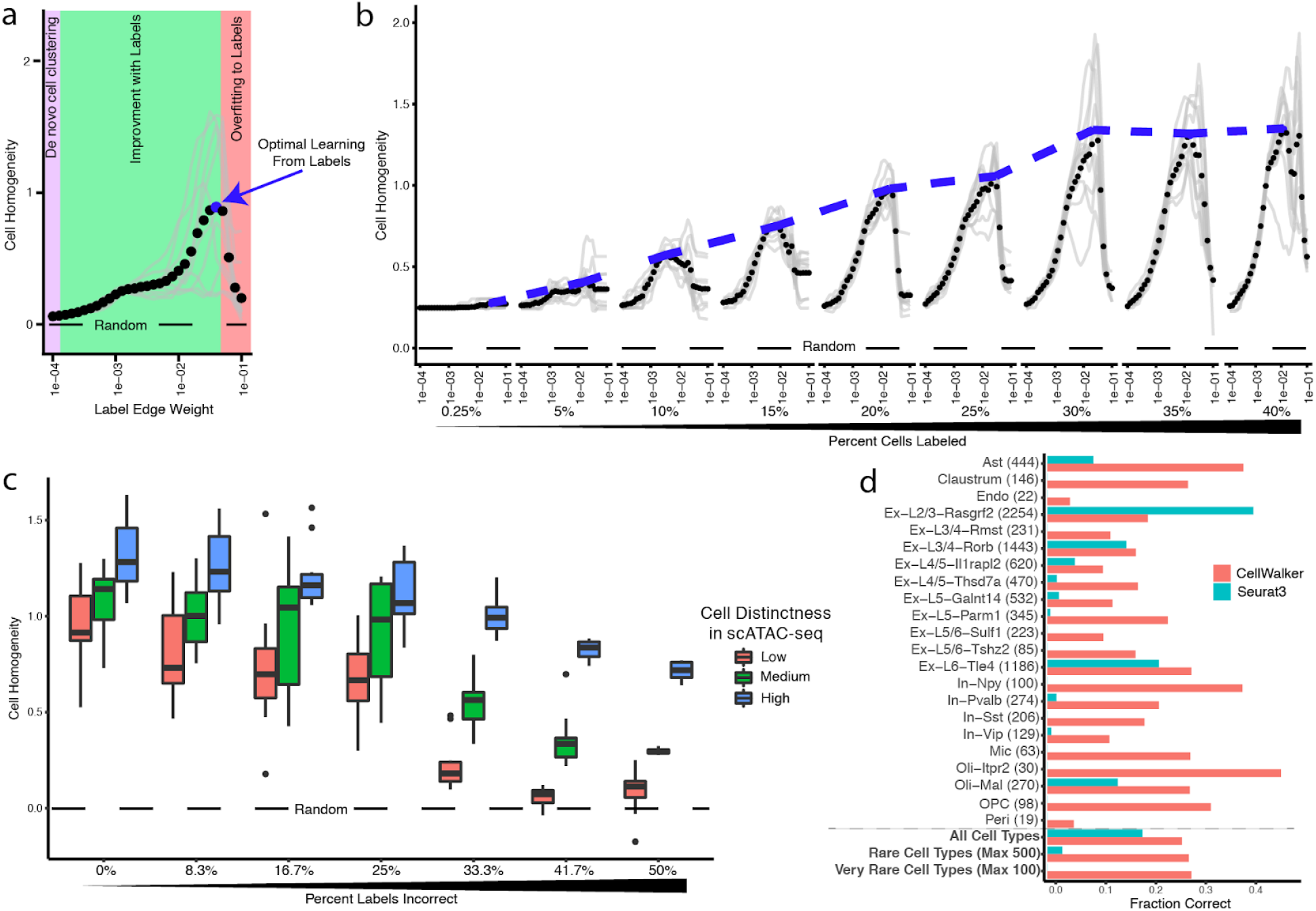
CellWalker Correctly Labels Cells in Simulations and Validation Data. **a.** Label-edge weight, defined as the ratio of label-to-cell edges versus cell-to-cell edges, (x-axis) is tuned in order to optimize cell homogeneity, a measure of the separability of cells of different types (y-axis). When edge weight is low the output is more similar to de-novo cell clustering (purple area) and when it is high the output becomes more similar to directly assigning labels to each cell (red area). Higher values of cell homogeneity indicate improved ability to distinguish between cells of different types, while a cell homogeneity of 0 is equivalent to no difference between within-cell-type and between-cell-type influence (dashed line). Black dots indicate mean performance across ten simulations (gray lines) **b.** As the percent of cells with labeling edges increases (x-axis) optimal cell homogeneity does as well, up to 30& labeled (purple line). **c.** The distribution of peak cell homogeneity scores across simulations when cell distinctness in scATAC-seq is low, medium, and high. As a higher percent of labels is incorrect (x-axis) performance begins to decline, particularly when initial cell distinctness is low. **d.** CellWalker correctly labels cells from the ATAC portion SNARE-seq data (number of cells of each type in parenthesis) with no dropoff for rare (max 500 cells) and very rare (max 100 cells) cell types.

Next, we tested CellWalker on mouse cortex SNARE-seq data which includes both scRNA-seq and scATAC-seq reads for each cell^20^. We analyzed the scATAC-seq portion of the data with CellWalker. For cell type labeling, we integrated the scATAC-seq data with differentially expressed marker genes previously derived from clustering the scRNA-seq portion of the SNARE-seq data. Performance was evaluated using the held-out scRNA-seq label for each cell that was identified in the original publication. We tuned the edge weight parameter to optimize cell homogeneity (as in our simulations) and observed that this closely mirrors optimization of the fraction of cells labeled correctly, validating cell homogeneity as a measure of how well cell types are resolved (Supplemental Figure 3). We compared CellWalker to label transfer, as implemented in Seurat^15^, and found that CellWalker labels more cells correctly (Figure 2d). This advantage is greater when considering only rare cell types and very rare cell types (Figure 2d, bottom). Taken together, these analyses of SNARE-seq data indicate that integration of label data provides a substantial advantage towards resolving cell types in scATAC-seq data.

### Identification of cell types in the developing brain

Given the ability of CellWalker to identify rare cell types in brain SNARE-seq data, we next applied it to a scATAC-seq study of the human telencephalon with multiple biological replicates spanning mid-gestation^19^. Previous work generated a cell type atlas in similar samples based on extensive analysis of scRNA-seq data^17^. Using this atlas as external labeling data, we used CellWalker to compute a full influence matrix across all these labels and 30,000 scATAC-seq cells. First, using the label-to-label portion of the influence matrix, we hierarchically clustered all labels and observed high agreement with clustering based on scRNA-data (Figure 3a). Next, using label-to-cell influence we scored each cell assigning it the highest scoring label. This produces a “fuzzy” labeling of cells, representing the fact that a scATAC-seq cell may be strongly connected through the network to multiple cell types. Nonetheless, we observed that most cells belong strongly to one type, indicating that most transcriptional states observed in scRNA-seq are associated with a distinct open chromatin signature in scATAC-seq (Figure 3b).

**Figure 3.**
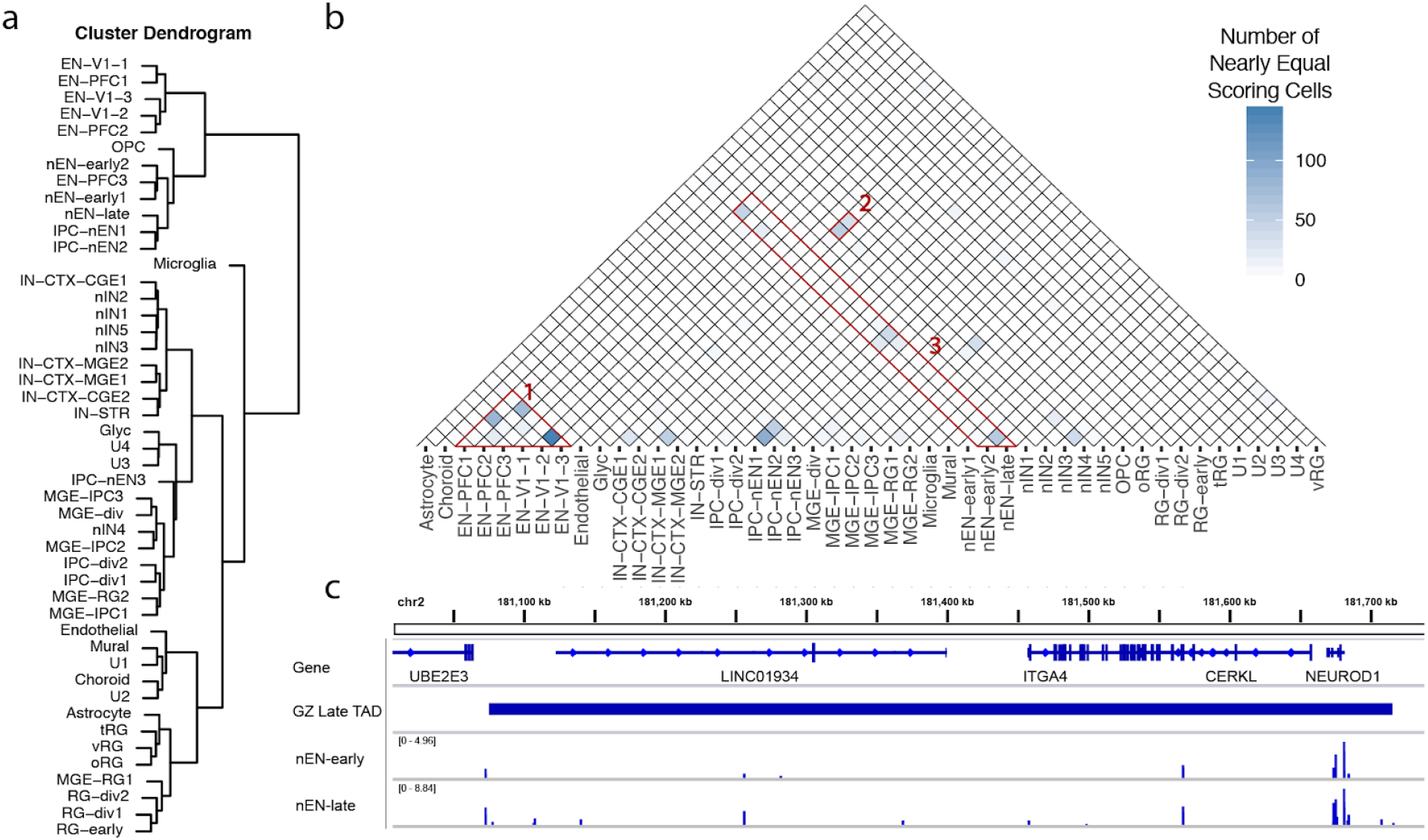
CellWalker Identifies Cell Types in the Developing Brain. **a.** Hierarchical clustering of label-to-label influence identifies cell types similar in scATAC-seq. **b.** A portion of cells has two nearly equal labels. Some are very similar mature lineages (group 1) while others correspond to progressions of neuronal development (groups 2 and 3). **c.** The region surrounding the late TAD containing the neurodevelopment transcription factor *NEUROD1*. Throughout this TAD accessibility outside the gene body and promoter increases between early and late newborn excitatory neurons and is highly correlated with excitatory neuron progression score (spearman correlation coefficient 0.56).

In a few cases we observed groups of cells with multiple nearly equally scoring labels, indicating intermediate membership in multiple cell types and revealing transcriptional states that correspond to highly similar open chromatin profiles. Some of these relationships, such as visual cortex (V1) and prefrontal cortex (PFC) excitatory neurons, represent similar types of maturing neurons that are present in two brain regions (Figure 3b, group 1). Others correspond to progressions of neuronal development. For example, the newborn interneuron and caudal ganglionic eminence (CGE) cortical interneuron cell types have shared influence on a large group of scATAC-seq cells (Figure 3b, group 2). Similarly, we observe cells that score highly as combinations of intermediate progenitor cells, early newborn excitatory neurons, late newborn excitatory neurons, and maturing excitatory neurons (Figure 3b, group 3), suggesting that these transcriptional states cannot be resolved in scATAC-seq.

To explore whether these indeterminate cell types represent limitations of scATAC-seq data, failures of the CellWalker model, or cases where transcription changes without large changes in open chromatin, we took a closer look at early and late newborn excitatory neurons. These are fairly large, identifiable cell types in scRNA-seq^17^. We assigned each scATAC-seq cell an excitatory neuron progression score based on the difference between the influence of the early and late newborn excitatory neuron labels. Therefore, a higher excitatory neuron progression score indicates a later newborn excitatory neuron. Using this score, we observe that while there is a small distinct set of early newborn excitatory neurons, the majority of newborn excitatory neurons fall evenly between the two types with many scores near zero (Supplemental Figure 4a). This indicates that there is a continuous gradient of changes in chromatin accessibility rather than large-scale difference between transitioning cell types. This appears to be a biological difference between the dynamics of gene expression versus open chromatin during developmental transitions, though higher coverage scATAC-seq data could potentially alter this conclusion.

### Cell-type specific annotation of loci

We next sought to determine if cell type annotations could be used to characterize the biology of loci based on chromatin accessibility at distal regulatory elements. Because most distal regulation occurs within Topologically Associated Domains (TADs)^21^, we sought to determine if the transition from early to late excitatory neurons could be attributed to differences in TADs between cell types. It is generally believed that Hi-C contact maps derived from bulk data represent the average of a mixture of cells^22^. We correlated the distal accessibility (defined as outside a gene body or promoter) of TADs derived from the germinal zone (GZ) of the mid-gestation developing human cerebral cortex^21^ with excitatory newborn progression score and found that the distribution of correlations is significantly bimodal (empirical *p*-value = 0.021). This means that the accessibility of GZ TADs distinctly either correlates or anti-correlates with cell state progression from early to late excitatory neuron. As a control, we find that the median distance of peaks to genes and the number of peaks per TAD do not correlate with excitatory neuron progression (Supplemental Figures 4b and 4c). We therefore classified GZ TADs as early or late depending on their correlation with excitatory neuron progression. As a validation of the classification of these TADs, we find that the expression of genes in early TADs negatively correlates with excitatory neuron progression score, while the expression genes in late TADs correlates positively (median correlations of −0.62 and 0.22 respectively). Thus, subtle changes in chromatin accessibility between early and late newborn excitatory neurons may be associated with cell-type specific TADs. The ability to separate TADs by cell type enables a greater understanding of gene regulation in complex tissues such as the human brain. A similar strategy could be applied to other annotations of loci, such as linkage disequilibrium (LD) blocks or expression quantitative trait loci (eQTLs).

Several key genes involved in neuronal development lie in early or late TADs indicating their expression may be distally regulated. Notably, the neurogenic differentiation gene *NEUROD1* lies in a late TAD with higher levels of accessibility late than early throughout the TAD but similar accessibility in the gene body and promoter (Figure 3d). Correspondingly, *NEUROD1* has two-fold higher mean transcripts in late than early newborn excitatory neurons (mean 73 TPM early vs 131 late). This indicates that the gene expression differences of *NEUROD1* are potentially driven by distal enhancers. Conversely, *TENM4*, which is involved in establishing neuronal connectivity during development^23^, lies in an early TAD and is less expressed in late newborn excitatory neurons (mean 350 TPM early vs 249 late). Deciphering the cell-type specific regulation of these genes is an important step towards understanding how differences in genotype lead to their misexpression and linked diseases.

### Cell-type specific annotation of enhancers

It is generally believed that many enhancers involved in brain development function in a cell type specific manner^19^. CellWalker provides a way to explore this idea. We mapped enhancers derived from bulk ATAC-seq on microdissected tissue across the mid-gestation human telencephalon^18^ to cell types based on cell-to-label influence (Figure 4a). As expected, we find that the many enhancers specific to the ganglionic eminence map to intermediate progenitor cell types, while enhancers in other regions primarily map to types of excitatory neurons^17^. As further validation, we also observe that enhancers from cell types labeled to specific regions such as the medial ganglionic eminence (MGE) map to regions sampled from the ganglionic eminence (Supplemental Figure 5a). These findings demonstrate that cell types resolved in scATAC-seq data with CellWalker can be used to annotate enhancers discovered in bulk ATAC-seq. This strategy combines the benefits of high coverage in bulk data with the cell-type information in scATAC-seq.

**Figure 4.**
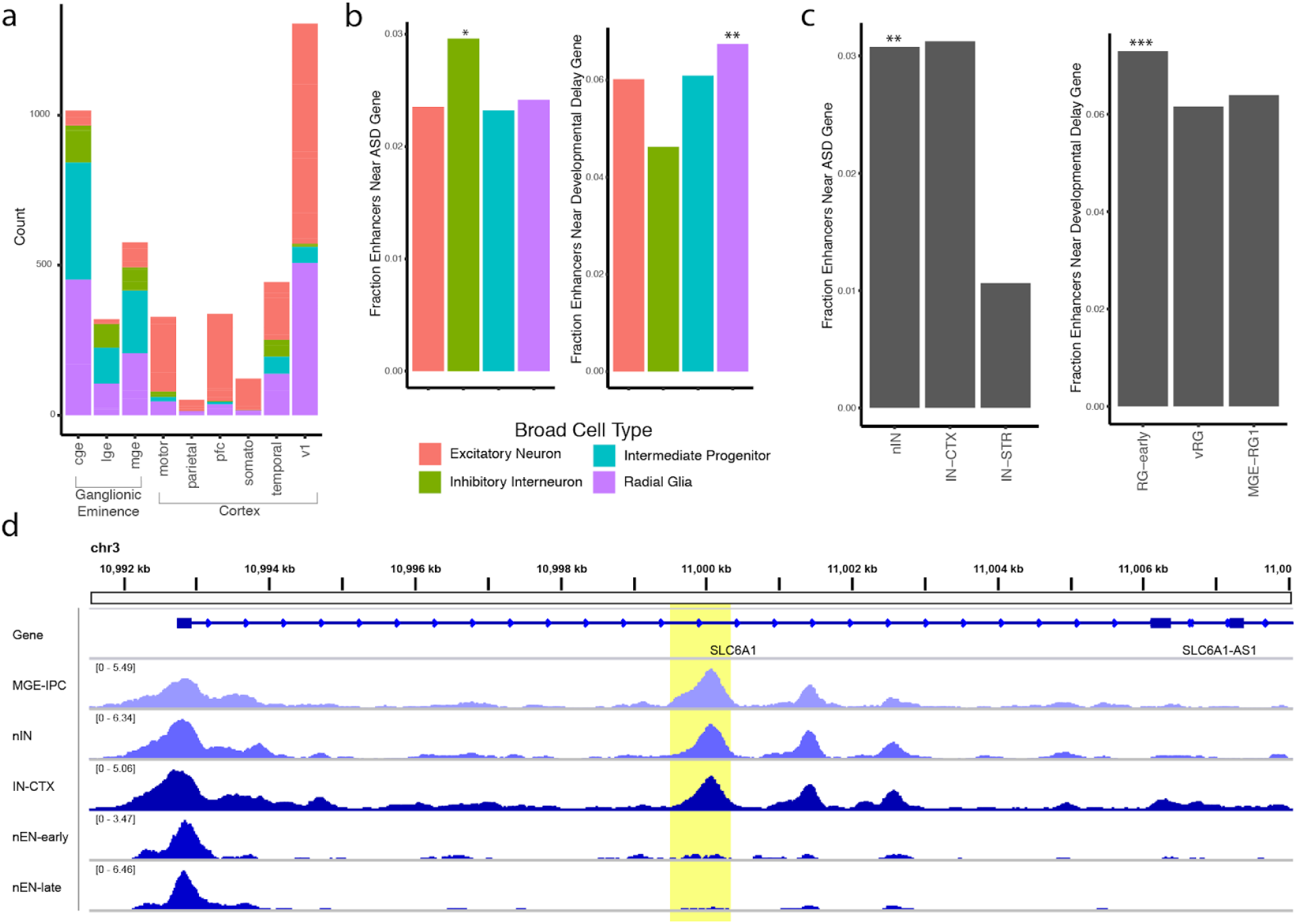
Cell-type Specific Annotation of Enhancers. **a.** CellWalker maps enhancers generated from microdissected brain regions to cell types using cell-to-label influence. **b.** A significant fraction of inhibitory interneuron specific enhancers are closest to ASD related genes (left) and a significant fraction of radial glia specific enhancers are closest to developmental delay related genes (left). **c.** For ASD of the interneuron subtypes, newborn interneurons (nIN) and cortical interneurons (IN-CTX) are enriched, but there are only enough newborn interneuron specific enhancers to achieve statistical significance (left). For developmental delay genes, early radial glia (RG-early) are enriched (left). **d.**The region surrounding an intronic SLC6A1 enhancer (yellow highlight). The enhancer is accessible throughout interneurons, but not in excitatory neurons. (* represents significance at FDR < 0.1, ** at FDR < 0.05, and *** at FDR < 0.01)

We next examined whether cell-type specific enhancers are associated with disease genes. First, we considered sets of genes near significant variants detected in a collection of Genome-Wide Association Studies (GWAS)^24^. Testing for associations with enhancers active in four broad cell types, among these gene sets we found significant enrichment for enhancers near genes associated with a collection of neurological diseases as well as many measures for developmental delay (significant at FDR < 0.1, Supplemental Table 1). We therefore decided to take a closer look at curated lists of genes linked to Autism Spectrum Disorders (ASD) and developmental delay^25^. We found that enhancers specific to radial glia are significantly associated with genes linked to developmental delay (Figure 4b, right), among which enhancers specific to early radial glia are significant (Figure 4c, right). Enhancers specific to inhibitory interneurons are significantly associated with genes linked to ASD, in agreement with previous studies^26,27^ (Figure 4b, left). Among these, enrichment was significant for newborn interneurons (Figure 4c, left). Recently, an enhancer of *SLC6A1* which is accessible in cells in the ganglionic eminence has been linked to ASD^18^. We found that this enhancer maps to both intermediate progenitor cells located in the ganglionic eminence as well as to newborn interneurons, and is accessible in cells predicted to belong to these cell types (Figure 4d). As validation, we find that this peak is accessible in intermediate progenitors and interneurons but not excitatory neurons in ATAC-seq data on FACS sorted cells^28^ (Supplemental Figure 5c). Interestingly, while the enhancer is most strongly linked to early stages of interneuron development, expression of *SLC6A1* increases throughout the course of interneuron development (Supplemental Figure 5b). It is possible therefore that *de novo* mutations observed in this enhancer in ASD individuals contribute to changes in the initiation of *SLC6A1* expression, which could influence the timing of interneuron development. This strategy for determining cell-type specific effects can be applied to other loci to better understand their potential roles in disease and cell differentiation.

## Discussion

The development of high-throughput sequencing technologies has enabled an explosion of data generation, necessitating techniques to integrate these data into knowledge and testable hypotheses. Using CellWalker, we were able to uncover cell-type specific signals based on a combination of bulk and single cell data. This was made possible with external data about known cell types which helped overcome the low signal-to-noise ratio present in scATAC-seq data. This strategy is broadly applicable as there are already vast amounts of bulk RNA-seq, bulk epigenomics, scRNA-seq, and other cell atlas data for tissues, organoids, and cell lines related to samples where scATAC-seq is being performed. Here, we applied CellWalker to neurodevelopment. In another study, we used CellWalker to map transcriptional disease states from mouse heart data to scATAC-seq data in matched tissues and uncovered cell-type specific enhancers activated by stress. CellWalker is computationally efficient enough to enable even larger scale integrations of such data (Supplemental Figure 5).

CellWalker extends naturally to incorporate multiple cell atlases simultaneously. This presents a myriad of possible opportunities such as measuring the influence between disease and non-disease labels for similar cells. For example, an atlas of the developing brain could be used together with cell types derived from post-mortem brains of individuals with ASD to directly measure the relationships between those sources of labeling data. An alternative possibility is to use CellWalker to transfer labels across species. Alternative integrations of data are possible by, for example, using the each labeled multiple times, but weighing edges differently based on bulk measurements of histone markers. Much of the power of CellWalker lies in its generalizable network model.

The cellular complexity of the developing human brain presented an ideal testbed for CellWalker. We were able to detect rare cell types and tease out distal regulatory programs by integrating bulk data with scATAC-seq data. However, as more single-cell data is generated with higher read depths and greater cell coverage it may turn out that our data simply did not have the power to uncover the true underlying regulatory landscape. Rare intermediate cell types that correspond better with identified transcriptional states may exist. Furthermore, new data that simultaneously measures multiple epigenetic and transcriptional attributes in the same cell will soon begin to enable the detection of cell-specific links between regulation and expression^29^. However, while these technologies continue to be developed and improved, exploiting existing troves of bulk data provides a powerful avenue towards understanding cell-type specific regulation.

## Methods

### CellWalker

The network constructed by CellWalker consists of two types of edges: cell-to-cell and label-to-cell. The former has an edge weight corresponding to the similarity between pairs of cells in scATAC-seq data. The latter has an edge weight corresponding to the similarity between the given labeling feature and each cell. With this general approach, it is possible to add a large variety of external data to the model. Although these edges may be sparsely connected to cells, the edges between cells distribute information. CellWalker includes a single parameter, label edge weight, which determines the ratio of the weight of label-to-cell edges relative to the weight of cell-to-cell edges. To diffuse the information from all data sources across the network, we implemented a random walk with restarts. A unit amount of information is initialized at each node. Then at each time step, a fixed portion restarts and the remainder propagates across each edge connected to the node, proportionally to edge weights. Even cells poorly annotated with external data will receive information about those annotations via cells that are similar. This algorithm is equivalent to an insulated heat diffusion graph kernel. To implement diffusion, we first compute a q-by-q walk matrix *W* encoding the fraction of information that must move to each neighboring node in each time step, where q is equal to the total number of nodes in the graph. This is 0 if the nodes have no edge between them and the fraction of total weight of edges for each node otherwise. In matrix notation the computation is *W=D*^−1^*A*, where *D* is a diagonal matrix of the sums of edge weights for each node and *A* is the adjacency matrix representing the graph. Given this formulation of the walk matrix *W* and a non-zero restart probability α, the walk always converges to a stationary distribution. Due to this property, there is a closed form solution for the q-by-q influence matrix *F*, which defines the amount of information that reaches each node from each other node and is computed as *F* = α(*I* ­ (1 ­ α)*W*)^−1^ . Prior work has examined how different settings of alpha distribute information to neighboring nodes and found that a restart probability between 0.4 and 0.6 encodes graph structure well with only minor variance in information in that range^30^. Based on this, we set our restart probability to 0.5.

### Simulations

We generated artificial cells that emulate high quality scATAC-seq processed by the SnapATAC^12^ pipeline as follows. For each of n cells we sampled the number of total reads for that cell from the distribution of reads per cell we observed in real data (median 5,500 reads per cell). We then distributed those reads across p bins proportionally to the distribution of reads per bin observed in real data. This resulted in a p-by-n count matrix of n cells and p bins. We split the pool of generated cells into two cell types and gave the cells low, medium, or high within-type distinctness by splitting bins evenly across cell types and adding a fixed percent (1, 5 or 10 respectively) of additional reads across those bins to each cell. In order to label cells, we generated two label nodes and created edges from these nodes to cells with a weight of 1 depending on the simulation scenario. Cell-cell edges were given a weight of the Jaccard similarity of each cell's bins. For each simulation we ran CellWalker 10 different assignments of cell-label edges for each of a range of label edge weights between 10^−4^ and 10^−1^ for 400 cells of each cell type. We evaluated the ability of CellWalker to seperate the two cell types using cell homogeneity which we computed directly from the influence matrix *F* as the log of median ratio of information between cells within the same cell type to information between cells of different cell types. To test the importance of label-to-cell edges we tested labeling between a single cell in each cell type up to labeling 40% of cells using medium cell distinctness. We tested the importance of cell mis-labeling by labeling 15% of cells correctly and adjusting the number of mislabeled cells between a single cell and 15% of cells (not necessarily mutually exclusively). In tests for robustness to noisy reads we randomly added or removed a fixed percentage of all reads in each cell. Finally, to test if CellWalker is able to distribute labels to cell populations even without any initial labeling, we generated an additional set of 400 cells with no labeling edges. Rather than give these cells medium, low, or high within group distinctness, they were made more similar to one of the previous cell types by being generated by randomly sampling reads proportionally to bins from either of the other two cell types, with the proportion of bins from the cell type adjusted between 10 and 50 percent. We additionally used simulated data to determine how CellWalker’s runtime and memory usage scales with the number of non-zero bins in the cell-by-bin matrix and found that both relationships are linear (Supplemental Figure 2).

### Data Processing and Analysis

We downloaded the cell-by-peak matrix for the scATAC-seq portion and the cell-by-gene matrix for the scRNA-seq portion of the SNARE-seq data for the adult mouse cerebral cortex^20^. We additionally downloaded the cell type labels assigned to each cell as well as marker genes for each cell type, which includes the log fold change of expression for each marker in the given cell type compared to other cells. We ran CellWalker on this data by computing the Jaccard similarity between binarized peak accessibility vectors for cells for cell-to-cell edges and the fraction of each cells peaks that are in marker’s gene body or promoter (2kb upstream of TSS) for a given cell type scaled by the log-fold change in expression of each marker for label-to-cell edges. We tested label edge weights between 10-2 and 10^4^ and computed both the cell homogeneity and the fraction of exact label matches at each weight (Supplemental Figure 3). We found that the two follow nearly identical trends implying that cell homogeneity is a good proxy for correct labeling. For comparison, we ran Seurat3^15^ on the cell-by-peak and cell-by-gene matrices and assigned labels using default parameters for anchor transfer between the two datasets.

Multi-sample mid-gestation human telencephalon scATAC-seq data from psychENCODE was previously processed using SnapATAC to generate a large cell-by-bin matrix^19^ and a previously derived set of marker genes was used for labeling^17^. As before, cell-to-cell edge weights were computed using Jaccard similarity and label-to-cell edge weights were computed as the sum of normalized SnapATAC generated gene accessibility score for each marker scaled by that marker genes log-fold change in expression. We tested label edge weights between 10^−2^ and 10^4^ and selected a weight of 1 as optimal. We hierarchically clustered labelling nodes using the euclidean distance between label-to-label vectors and “hclust” with default parameters in R^31^. To compute cell label scores from label-to-cell influence we compute z-scores for each column and then rescale to a maximum score of 1. We considered a cell to have two nearly identical labeling scores if the top two highest scoring labels were within the bottom 10% of all differences between the two highest scoring labels for all cells.

To compute an excitatory neuron progression score for each cell we took the difference between the nEN-early2 and nEN-late score for each cell. For our analysis we only considered the subset of cells with nEN-early2 or nEN-late as their top label scores. 1,367 GZ TADS were previously generated^32^ based on HiC data taken from Won et al^21^. For each TAD we computed the fraction of each cell’s distal (non-promoter, not in gene body) peaks that were accessible in that TAD. Then for each TAD we correlated excitatory neuron progression scores for each cell with the distal accessibility of each cell. Negative correlations imply early active TADs and positive correlations imply late active TADs. To determine if this distribution of correlations is significantly bimodal we permuted TAD locations 1,000 times using the randomizeRegions function in regioneR^33^ (restricted to exclude blacklisted locations) and compared the median absolute values of correlations. Of the full set of TADs, we classified 290 as early TADs with correlation less than −0.5 and 247 as late TADs with correlation > 0.5. Mean transcripts (TPM) were computed from the same scRNA data that cell type marker genes were derived from^17^.

Regionally microdissected developmental brain enhancers were downloaded from Markenscoff-Papadimitriou et al^18^. To score each label for each enhancer we calculated the sum of cell-to-label influence across all cells which had a peak in the given enhancer. We split cell types into four primary labels following the groupings in Nowakowski et al^17^. GWAS data was downloaded from the NHGRI-EBI GWAS catalog^24^. A list of genes was generated for each disease or trait using all mapped or reported genes for each significant variant as annotated in the catalog. We omitted diseases and traits that had fewer than 100 associated genes. Disease gene sets were downloaded from Werling et al^25^. To estimate disease gene set enrichment, we computed an empirical *p*-value by comparing the fraction of enhancers that were closest to disease genes to the fraction closest to an equal size random sample of brain expressed genes, resampling 10,000 times. FDRs were computed using the Benjamini-Hochberg procedure. Cell tracks were generated using SnapATAC^12^. Peaks from FACS sorted cells were taken from Song et al^28^.

## Acknowledgements

GZ TADs were generated and made available by Sean Whalen^**32**^. Ryan Ziffra and Tom Nowakowski provided early access to processed scATAC-seq data and helpful feedback on the project^19^. Thank you to Kathleen Keough for suggestions to improve manuscript clarity and John Rubenstein for help interpreting our results in the context of neurodevelopmental gene regulation. This work was supported by research grants to K.S.P from: NIH/NIMH awards U01-MH116438, R01-MH109907, and R01-MH123178.

## Author Contributions

P.F.P. and K.S.P. designed the study, P.F.P. performed the analysis, P.F.P and K.S.P. wrote the manuscript. All authors read and approved the final manuscript.

