## Supplemental Figures for "CellWalker integrates single-cell and bulk data to resolve regulatory elements across cell types in complex tissues"

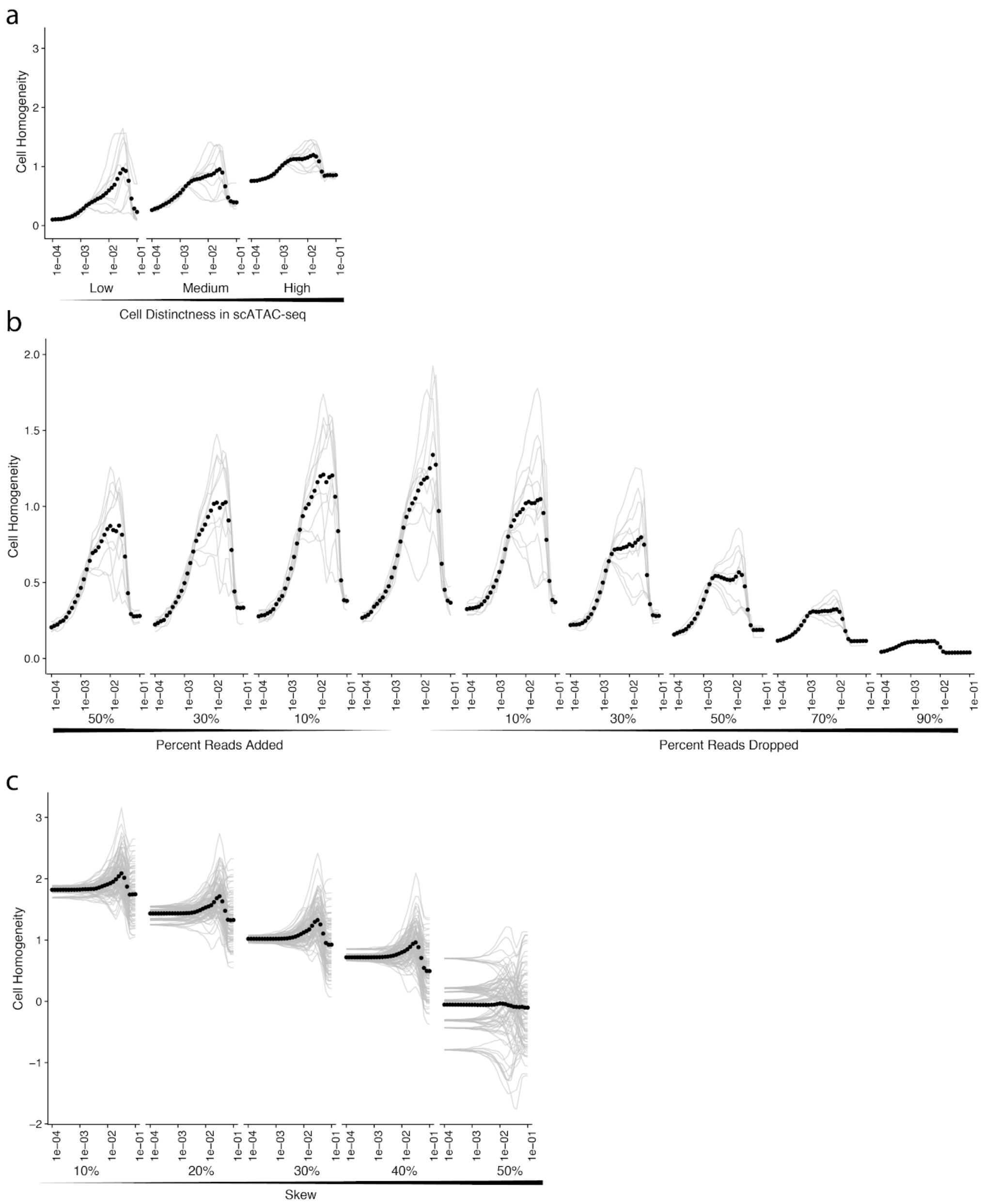


Supplemental Figure 1**. Additional Simulation Results. a.** As the within-type cell distinctness increases (x-axis) de-novo cell homogeneity increases a large amount (first point on each curve) but optimal cell homogeneity increases only slightly, indicating that within-cell type similarity can be low to achieve high cell homogeneity. **b.** As simulated reads are randomly added or removed (x-axs) cell homogeneity decreases. Adding random reads (left) only slightly decreases cell homogeneity. While removing reads (right) has a larger effect, even as many as 50% of reads can be dropped and optimal performance is still higher than the highest de-novo cell homogeneity. **c.** An unlabeled population of cells is less and less skewed towards one cell type (x-axis) with the proportion of bins from the cell type adjusted between 10 and 50 percent. Until the unlabeled cell type is exactly equally sampled from the other two types, cell homogeneity (in this case computed as the ratio of influence between the unlabeled cells and the two sets of labeled cells) remains high. Each level of skew includes 10 repeated random relabelings of 10 randomly generated sets of skewed cells.

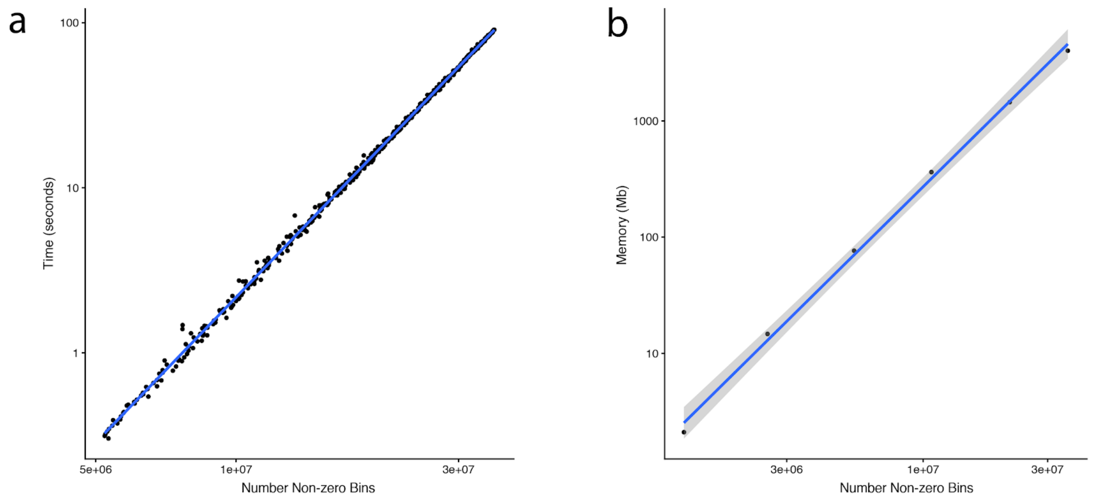


Supplemental Figure 2. **Runtime Analysis. a.** Running time for CellWalker (y-axis) versus the number of non-zero bins in the cell-by-bin matrix representing scATAC-seq data (x-axis). In our data, cells had a median of around 6,000 non-zero entries. We can extrapolate that it would take about 8 minutes to run CellWalker on a set of 10,000 cells. **b.** Memory usage of CellWalker versus the number of non-zero bins. We can extrapolate that it would take about 20Gb of RAM to run CellWalker on a set of 10,000 cells.

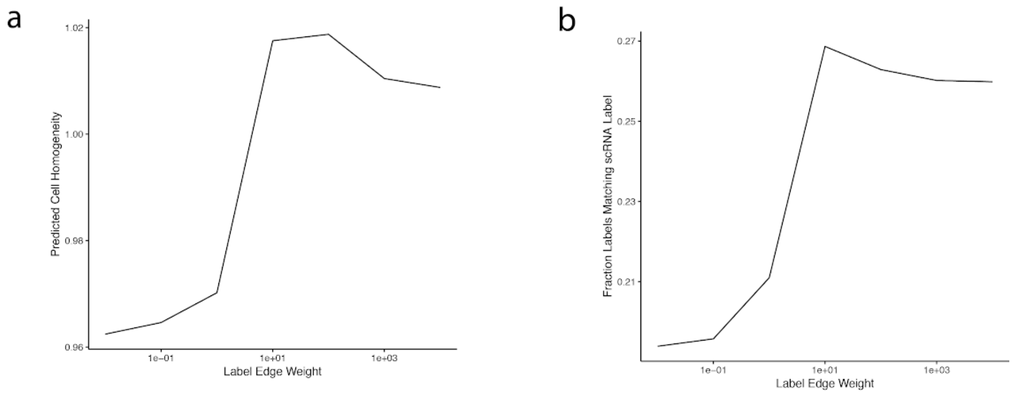


Supplemental Figure 3. **CellWalker Performance on SNARE-seq Data. a.** Cell homogeneity (y-axis) computed using the hidden scRNA-seq based cell labels as the median ratio of within-label vs out-of-label influence across possible settings of the label edge weight parameter (x-axis) **b.** Fraction of cells labeled the same as the hidden scRNA-seq based label (y-axis) across settings of the label edge weight parameter. The two measures peak near the same setting of the parameter.

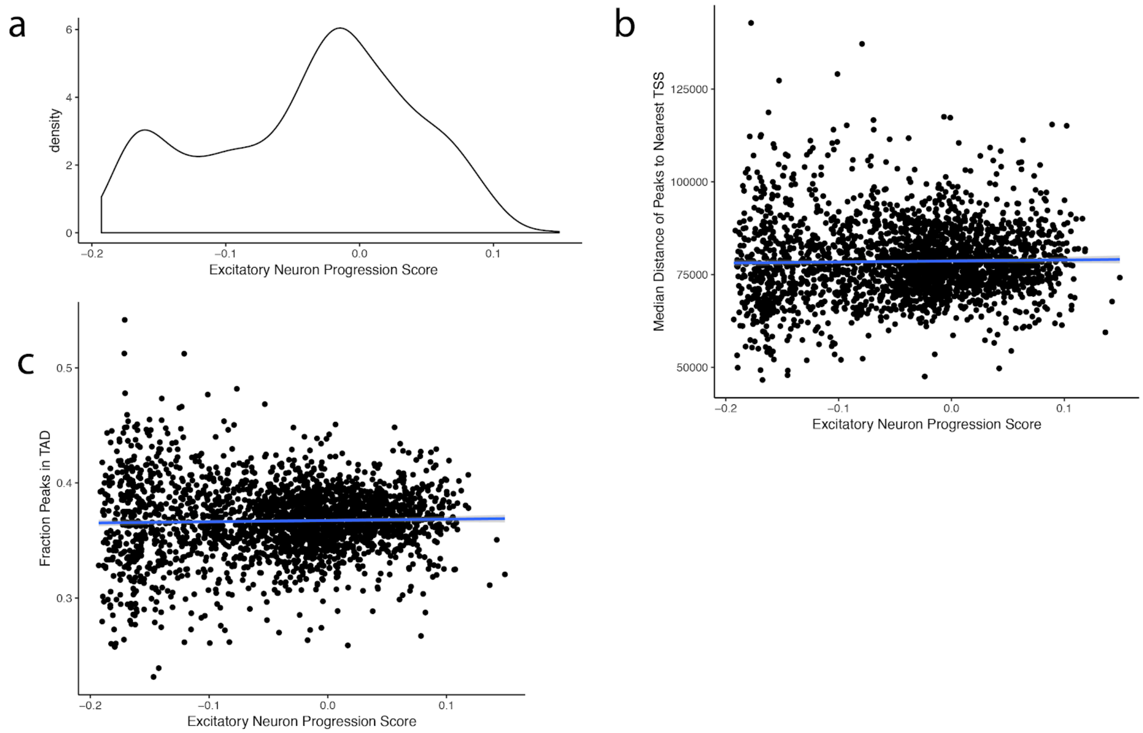


Supplemental Figure 4. **nEN Progression. a.** Distribution of excitatory neuron progression scores. **b.** Correlation between excitatory neuron progression score and the median distance from distal peaks to their nearest TSS. Each point is one cell, and the blue line is the best fit of a linear model. The two are not significantly correlated (Pearson's correlation coefficient 0.02, *p*-value=0.3) **c.** Correlation between excitatory neuron progression score and fraction of peaks in TADs. Each point is one cell, and the blue line is the best fit of a linear model. The two are not significantly correlated (Pearson's correlation coefficient 0.03, *p*-value=0.2).

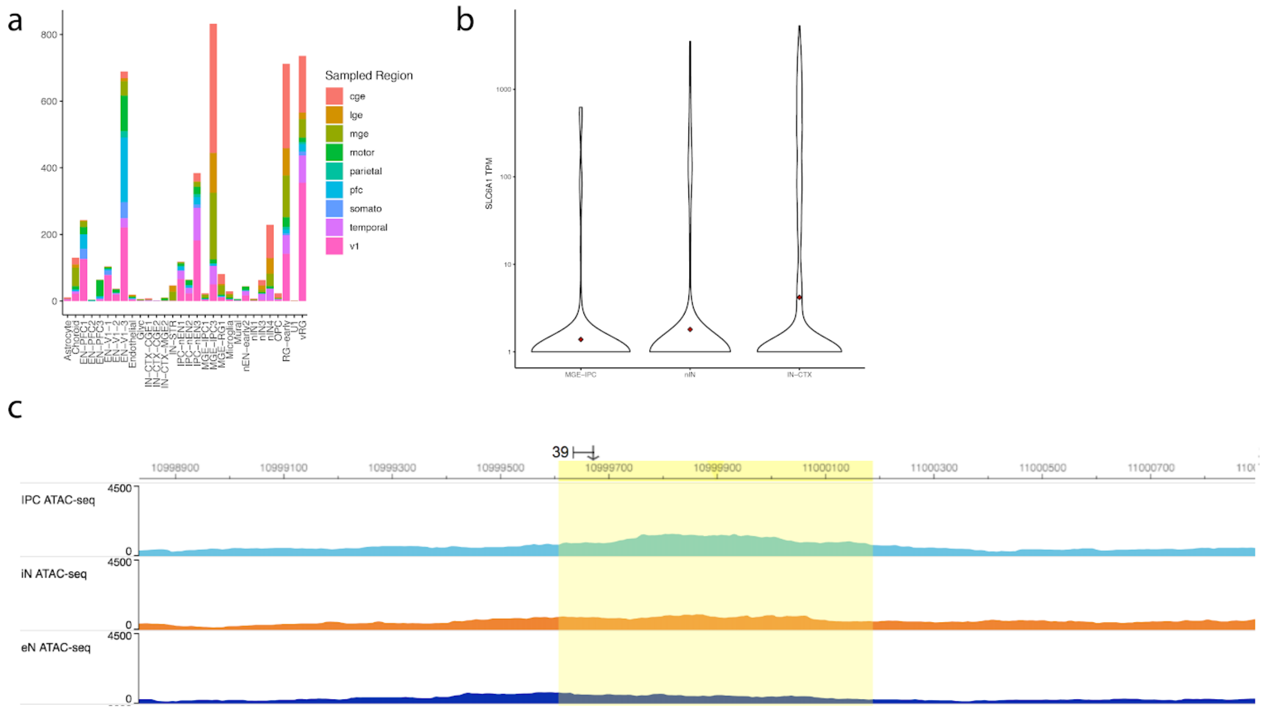


Supplemental Figure 5. **Cell-type Specific Enhancers. a.** The number of cell-type specific enhancers that uniquely map to enhancers from each microdissected region. **b.** TPM for SLC6A1 for each cell in scRNA-seq data with mean shown in red. **c.** Peaks in bulk ATAC-seq data for FACS sorted cells in the SLC6A1 enhancer (yellow highlighted region).
